## Supplementary Information for "Cell-free prototyping enables implementation of optimized reverse β-oxidation pathways in heterotrophic and autotrophic bacteria"

**Supplementary Figures**

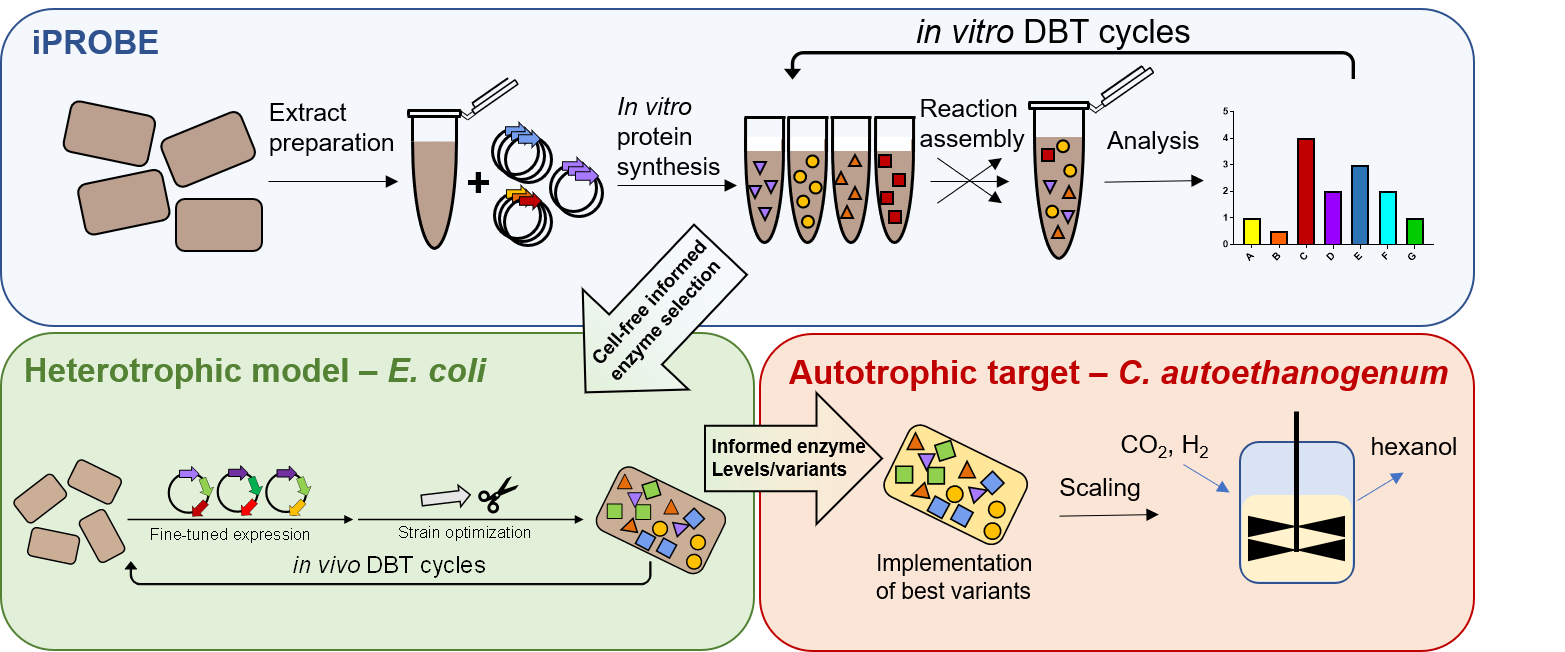

**Figure S1.** Scheme of the overall workflow for the project. Information gained in each system benefits all the other systems. iPROBE let to the identification of selective r-BOX variants. *E. coli* strain optimization led to the generation of r-BOX optimized CFPS extracts.

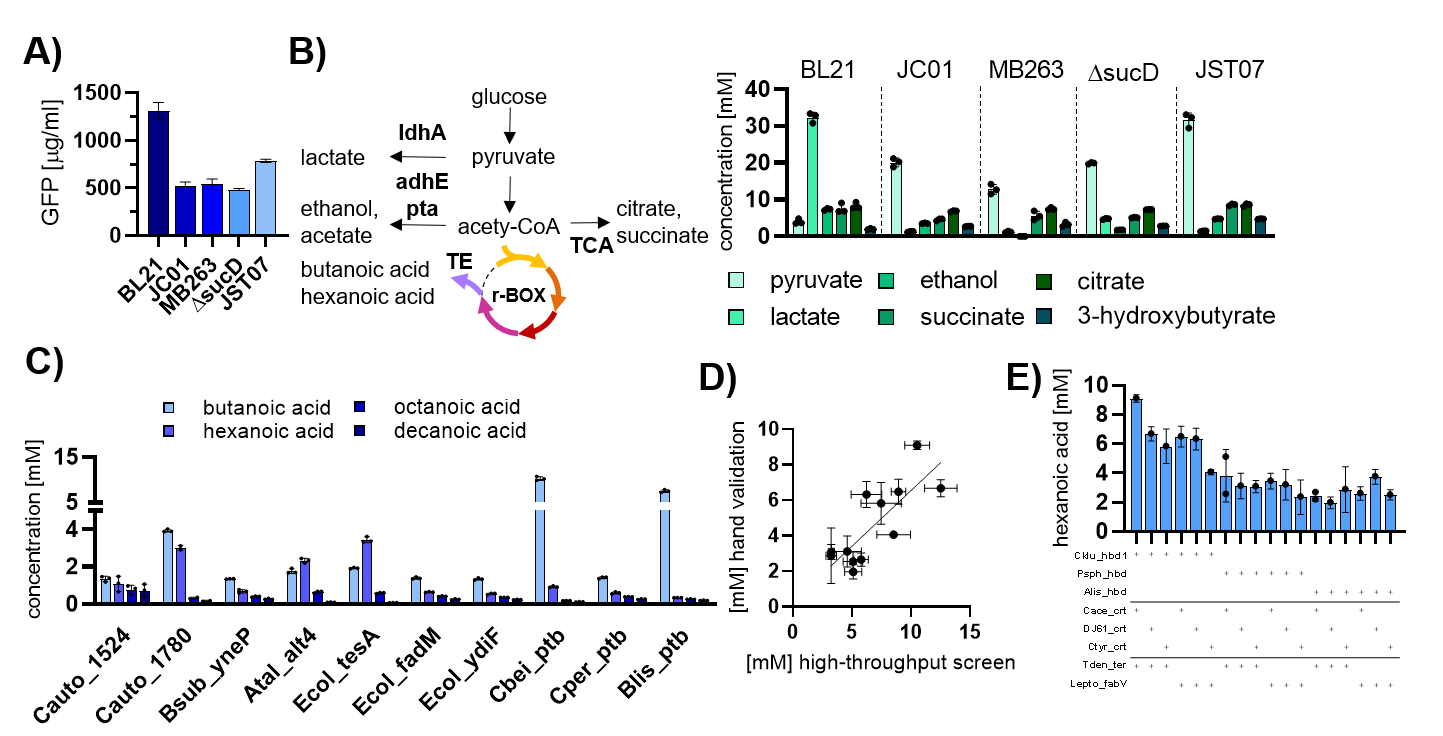

**Figure S2. (A)** Cell-free protein synthesis yields. **(B)** Side product quantification for all extracts using 0.3μM of Cn_bktB, Ck_hbd, Ca_crt, Td_ter and 0.15μM of Ec_tesA. **(C)** Screen of different thioesterases or a combination of a butyrate kinase (DJ052_buk) and phosphate butyryltransferases at 0.15μM keeping all core r-BOX enzymes the same as in S1B. Ec_tesA shows the highest specificity for hexanoic acid, Cb_ptb and Bl_ptb show high specificity for butanoic acid. **(D)** Select variants (top 15 combinations) measured both in the automated plate-based assay at 4μL volume and in **(E)** hand-pipetted assays at 30μL volume in Eppendorf tubes. The set shows a positive correlation with a slope of 0.63 ± 0.10 and an R^2^ of 0.59. Data represent n=3 independent experiments, with the standard deviation shown.

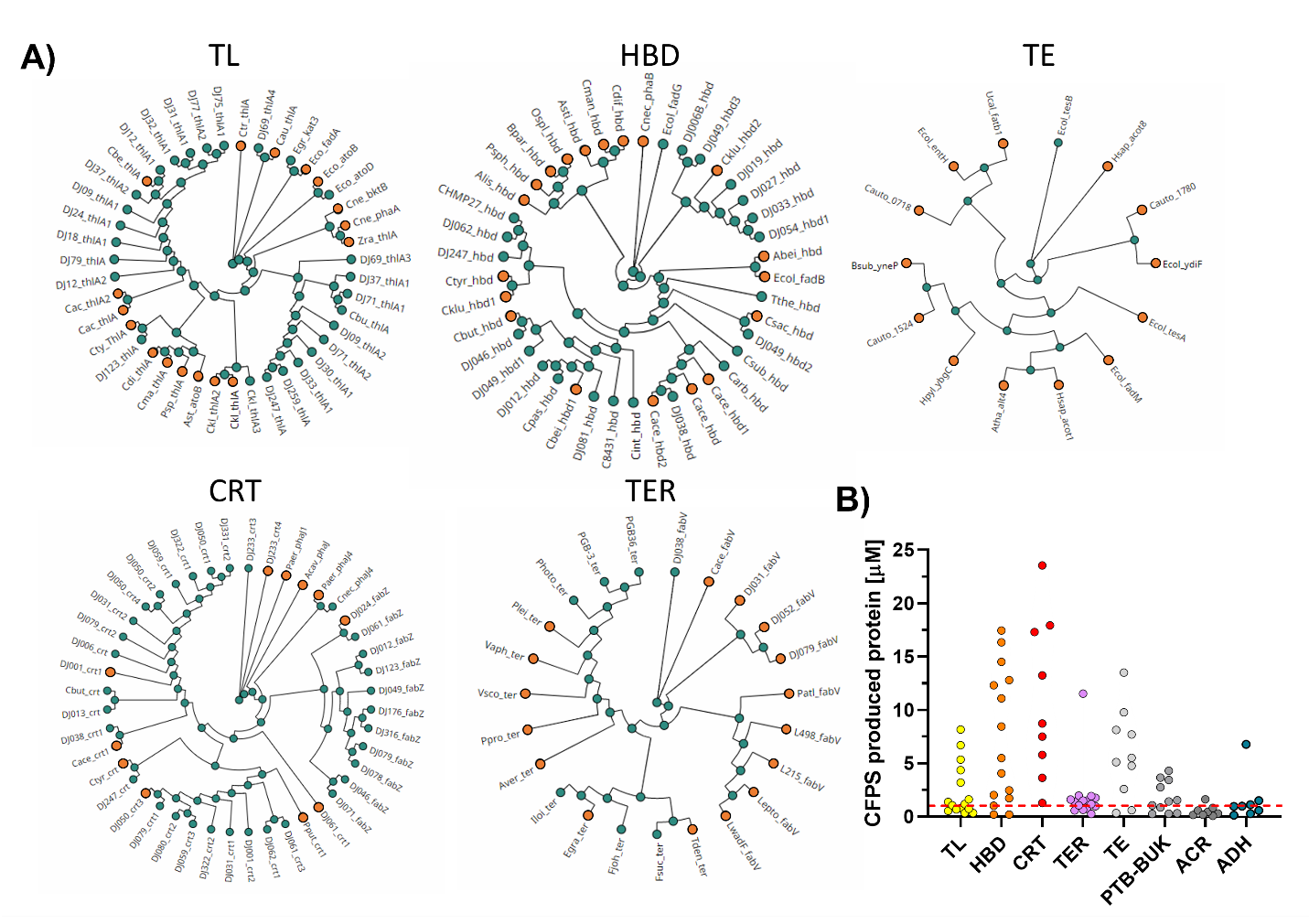

**Figure S3. (A)** Phylogenetic analysis of r-BOX enzymes using the MPI Tuebingen bioinformatics pipeline^1^. Orange dots highlight the proteins included in our cell-free analysis. TL = thiolase, HBD = hydroxybutyryl-CoA dehydrogenase, CRT = crotonase, TER = trans enoyl-reductase, TE = thioesterase. **(B)** Soluble protein yields for the 100 selected rBOX enzymes classed by function. Red line indicates the cut-off for soluble protein classification. Detailed list of enzymes and expression yields in supplementary sheet FigureS2_enzyme list.

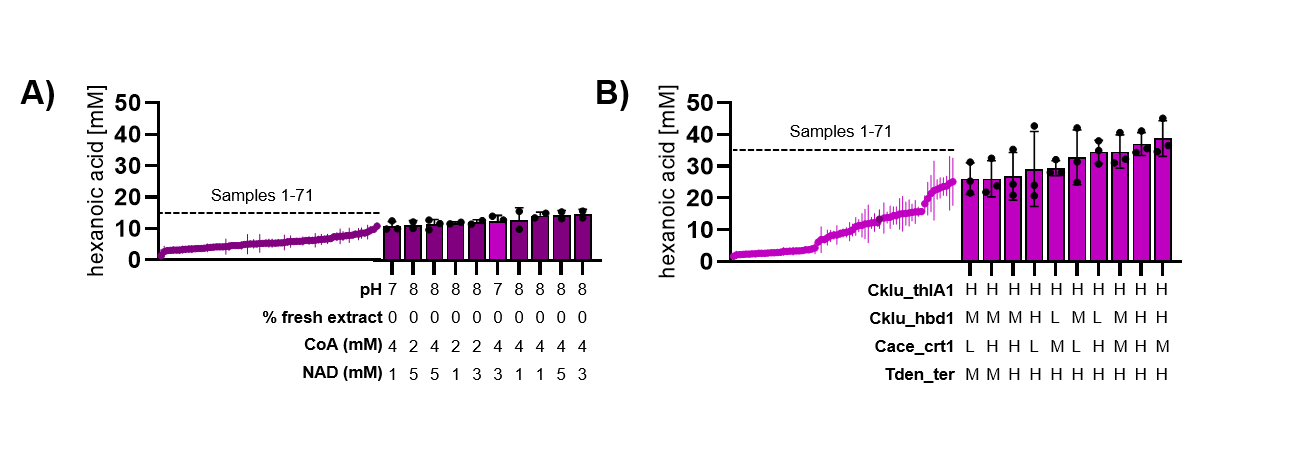

**Figure S4. (A)** Buffer, concentration of glycolytic enzymes and cofactor optimization for the best set of r-BOX enzymes from the screen. **(B)** Optimization of enzyme amounts by combinatorially testing all four enzymes at L = 0.05μM, M = 0.2μM and H = 0.6μM while keeping Ec_tesA at 0.10μM. Detailed results in supplementary sheet Figure2_screen. Data represent n=3 independent experiments, with the standard deviation shown.

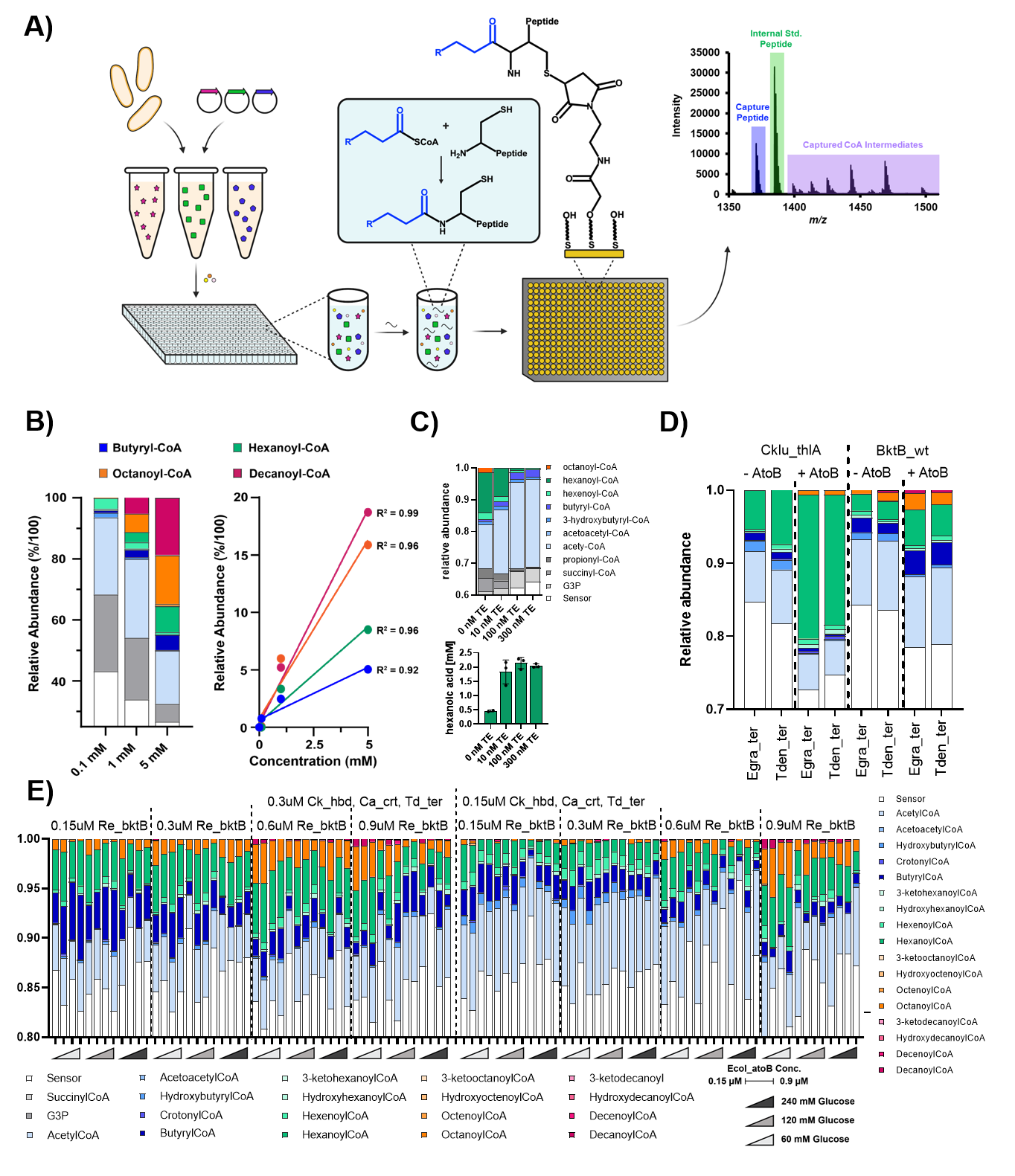

**Figure S5. (A)** General workflow for SAMDI-CoA measurements. CFPS-ME assays are assembled using automated acoustic liquid handling into 384 well plates. Samples are quenched and then derivatized via NCL to the capture peptide. Capture peptide is then bound to the SAMDI surface and measured via MALDI. **(B)** Empty spent CFPS-ME reactions were spiked with pure synthesized CoA-ester standards, processed and measured via MALDI. **(C)** Addition of termination enzyme (Ec_tesA) reduces the CoA-ester intermediates. Correlates well with amount of hexanoic acid produced as determined by GC-MS. All further SAMDI-CoA experiments were therefore done without addition of any termination enzyme. **(D)** Addition of a thiolase proficient at initializing r-BOX helped produce longer chain CoA-ester intermediates especially in the case of Ec_atoB and Re_bktB. **(E)** Optimizing *in vitro* conditions for build-up of long chain CoA-esters. Thiolase amount, addition of a second initializing thiolase Ec_atoB, as well as glucose input affected amount of >C6 CoA-ester formation. Core r-BOX enzymes did not seem rate limiting. All conditions were run and measured in quadruplicate (n = 4) and the average is displayed.
